# Ultrapotent miniproteins targeting the receptor-binding domain protect against SARS-CoV-2 infection and disease in mice

**DOI:** 10.1101/2021.03.01.433110

**Authors:** James Brett Case, Rita E. Chen, Longxing Cao, Baoling Ying, Emma S. Winkler, Inna Goreshnik, Swathi Shrihari, Natasha M. Kafai, Adam L. Bailey, Xuping Xie, Pei-Yong Shi, Rashmi Ravichandran, Lauren Carter, Lance Stewart, David Baker, Michael S. Diamond

## Abstract

Despite the introduction of public health measures and spike protein-based vaccines to mitigate the COVID-19 pandemic, SARS-CoV-2 infections and deaths continue to rise. Previously, we used a structural design approach to develop picomolar range miniproteins targeting the SARS-CoV-2 receptor binding domain. Here, we investigated the capacity of modified versions of one lead binder, LCB1, to protect against SARS-CoV-2-mediated lung disease in human ACE2-expressing transgenic mice. Systemic administration of LCB1-Fc reduced viral burden, diminished immune cell infiltration and inflammation, and completely prevented lung disease and pathology. A single intranasal dose of LCB1v1.3 reduced SARS-CoV-2 infection in the lung even when given as many as five days before or two days after virus inoculation. Importantly, LCB1v1.3 protected *in vivo* against a historical strain (WA1/2020), an emerging B.1.1.7 strain, and a strain encoding key E484K and N501Y spike protein substitutions. These data support development of LCB1v1.3 for prevention or treatment of SARS-CoV-2 infection.

## INTRODUCTION

Severe Acute Respiratory Syndrome Coronavirus 2 (SARS-CoV-2), the cause of the Coronavirus Disease 2019 (COVID-19) pandemic, has resulted in global disease, suffering, and economic hardship. Despite implementation of public health measures, SARS-CoV-2 transmission persists principally through human-to-human spread (Day, 2020; Li et al., 2020; Standl et al., 2020). SARS-CoV-2-induced clinical manifestations range from asymptomatic infection to severe pneumonia, multi-organ failure, and death. Although the underlying mechanisms that dictate disease severity are poorly understood, the immunocompromised, the elderly, and those with specific comorbidities (*e.g*., history of cardiovascular disease, diabetes, or obesity) are at increased risk for poor outcome (Zhou et al., 2020).

SARS-CoV-2 entry into target cells is facilitated by the spike glycoprotein through binding to its principal receptor, angiotensin converting enzyme 2 (ACE2) (Hoffmann et al., 2020; Letko et al., 2020). Once the virus is attached to the cell-surface, the spike protein is cleaved by the cell membrane-associated protease, TMPRSS2, resulting in membrane fusion and release of the viral RNA genome into the host cell cytoplasm (Hoffmann et al., 2020; Matsuyama et al., 2020). As the dominant antigen on the surface of the virion, the spike protein is the primary target of antibody-based countermeasures (Jeyanathan et al., 2020; Krammer, 2020). At present, a small number of antibody therapies and vaccines have been granted emergency use authorization (EUA) by the United States Food and Drug Administration to prevent or treat SARS-CoV-2 infection and disease. Nonetheless, viral evolution and the emergence of SARS-CoV-2 variants in the United Kingdom (B.1.1.7), South Africa (B.1.351), Brazil (B.1.1.248), and elsewhere jeopardize these countermeasures through potential loss-of-binding and diminished neutralization (Galloway et al., 2021; Leung et al., 2021; Tegally et al., 2020; Voloch et al., 2020).

We recently generated a panel of short, 56-amino acid miniproteins that bind the SARS-CoV-2 receptor-binding domain (RBD) with high affinity and potently neutralize authentic virus in cell culture with half-maximal effective concentration (EC_50_) values < 30 pM (Cao et al., 2020). Here, using a stringent model of SARS-CoV-2 disease pathogenesis in human ACE2 (hACE2)-expressing transgenic mice (Golden et al., 2020; Winkler et al., 2020a), we evaluated the efficacy *in vivo* of one of these miniprotein binders, LCB1. For our *in vivo* experiments, we evaluated two versions of LCB1: (a) an Fc-modified bivalent form, LCB1-hIgG-Fc9 (LCB1-Fc) that should extend half-life *in vivo* and engage effector arms of the immune system; and (b) a further optimized, monomeric form of LCB1 lacking an Fc domain, LCB1v1.3. Intraperitoneal administration of LCB1-Fc at one day pre- or post SARS-CoV-2 exposure conferred substantial protection including an absence of weight loss, reductions in viral burden approaching the limit of detection, and inhibition of lung inflammation and pathology. Intranasal delivery of LCB1v1.3 conferred protection as many as five days before or two days after SARS-CoV-2 inoculation. Dosing experiments revealed that LCB1v1.3 retained efficacy at pharmacologically attainable concentrations and was weakly immunogenic. Most importantly, LCB1v1.3 protected animals against the currently emerging B.1.1.7 United Kingdom variant and a SARS-CoV-2 strain encoding key spike substitutions E484K and N501Y present in both the South Africa (B.1.351) and Brazil (B. 1.1.248) variants of concern. Overall, these studies establish LCB1-Fc and LCB1v1.3 as possible treatments to prevent or mitigate SARS-CoV-2 disease.

## RESULTS

### LCB1v1.3 prophylaxis limits viral burden and clinical disease

Using computational design and functional screens, we previously identified LCB1 as a potent miniprotein inhibitor of SARS-CoV-2 infection (Cao et al., 2020). We modified LCB1 to generate two versions for *in vivo* testing: (a) we introduced polar mutations into LCB1 to increase expression yield and solubility without altering RBD binding (LCB1v1.3) and (b) we modified LCB1 by fusing it to a human IgG1 Fc domain (LCB1-Fc) to enhance bioavailability. LCB1v1.3 and LCB1-Fc bound avidly to a single RBD within the S trimer (**Fig 1A**) with dissociation constants (K_D_) of less than 625 and 156 pM, respectively (**Fig 1B**). LCB1v1.3 and LCB1-Fc also potently neutralized an authentic SARS-CoV-2 isolate (2019n-CoV/USA_WA1/2020 [WA1/2020]) (EC_50_ of 14.4 and 71.8 pM, respectively; **Fig 1C**).

**Figure 1.**
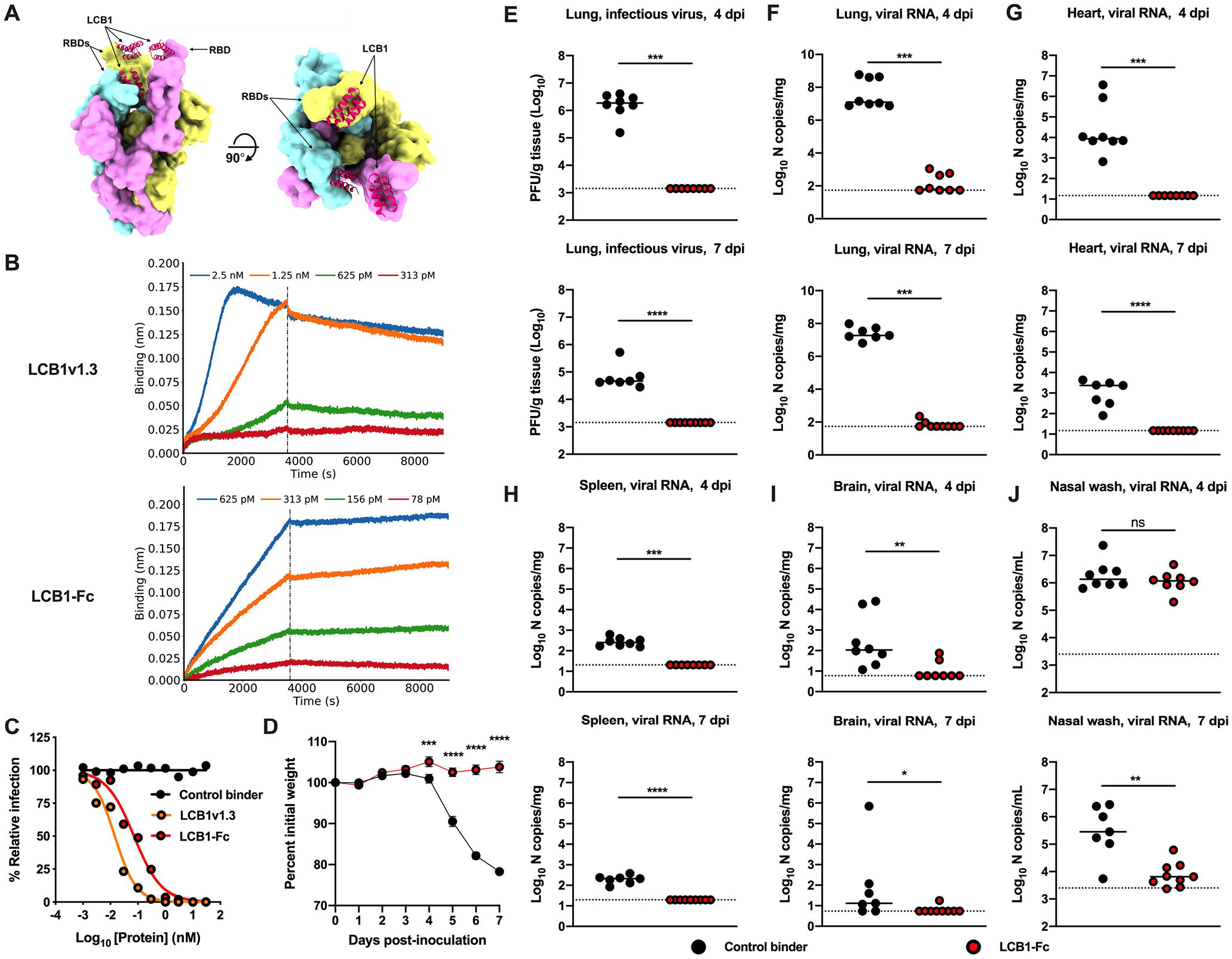
LCB1-Fc prophylaxis protects against SARS-CoV-2 infection. (**A**) Molecular surface representation of three LCB1v1.3 miniproteins bound to individual protomers of the SARS-CoV-2 spike protein trimer (left: side view; right: top view). (**B**) Binding curves of purified LCB1v1.3 and LCB1-Fc to SARS-CoV-2 RBD as monitored by biolayer interferometry (one experiment performed in technical duplicate). (**C**) Neutralization curves of LCB1v1.3, LCB1-Fc, or control binder against a SARS-CoV-2 WA1/2020 isolate (EC_50_ values: 14.4 pM, 71.8 pM, and >10,000 nM respectively; average of two experiments, each performed in duplicate). (**D-J**) 7 to 8-week-old female and male K18-hACE2 transgenic mice received 250 μg of LCB1-Fc or control binder by i.p. injection one day prior to i.n. inoculation with 10^3^ PFU of SARS-CoV-2. Tissues were collected at 4 and 7 dpi. (**D**) Weight change following LCB1-Fc administration (mean ± SEM; n = 8, two experiments: two-way ANOVA with Sidak’s post-test: *** *P* < 0.001, **** *P* < 0.0001). (**E**) Infectious virus measured by plaque assay at 4 or 7 dpi in the lung (n = 8, two experiments: Mann-Whitney test; *** *P* < 0.001). (**F-J**) Viral RNA levels at 4 or 7 dpi in the lung, heart, spleen, brain, or nasal wash (n = 8, two experiments: Mann-Whitney test: ns, not significant, * *P* < 0.05, ** *P* < 0.01, *** *P* < 0.001, **** *P* < 0.0001).

To determine the protective potential of these miniproteins against SARS-CoV-2, we utilized K18 human hACE2-expressing transgenic mice, which develop severe lung infection and disease after intranasal inoculation of SARS-CoV-2 (Golden et al., 2020; Winkler et al., 2020a). In prophylaxis studies, a single 250 μg (10 mg/kg) dose of LCB1-Fc administered by intraperitoneal injection (i.p.) one day prior to intranasal (i.n.) inoculation with 10^3^ PFU of SARS-CoV-2 WA1/2020 prevented weight loss compared to animals given a control protein (influenza A virus hemagglutinin minibinder) designed using similar computational methods (**Fig 1D**). After LCB1-Fc prophylaxis, infectious virus was not detected in the lungs at 4- or 7-days post-infection (dpi), whereas high levels were observed in animals administered control protein (**Fig 1E,** *top and bottom*). Similarly, viral RNA levels in the lung, heart, spleen, and brain of LCB1-Fc treated animals were at or near the limit of detection of the assay at 4 or 7 dpi (**Fig 1F-I**). LCB1-Fc treatment had no effect on viral RNA levels in nasal wash samples obtained at 4 dpi (**Fig 1J**), results that are similar to a recent study of a neutralizing human antibody in hamsters (Zhou et al., 2021). However, viral RNA levels were reduced at 7 dpi, suggesting that LCB1-Fc treatment accelerated viral clearance or prevented spread in the upper respiratory tract.

Diffuse alveolar damage, inflammation, and pneumonia are manifestations of COVID-19 lung disease, culminating in respiratory failure and a requirement for mechanical ventilation (Johnson et al., 2020; Kordzadeh-Kermani et al., 2020). We evaluated the capacity of LCB1-Fc to prevent the compromised lung function seen after SARS-CoV-2 infection of K18-hACE2 mice (Winkler et al., 2020a). At 7 dpi, mechanical ventilation tests of lung biomechanics in animals treated with LCB1-Fc showed no difference from naïve animals (**Fig 2A**), whereas mice receiving the control binder protein showed decreased inspiratory capacity and lung compliance as well as increased pulmonary resistance, elastance, and tissue damping, all consistent with compromised lung function. These biophysical properties resulted in disparate pressure-volume loops between control binder and LCB1-Fc treated or naïve animals. We also assessed the effect of LCB1-Fc treatment on SARS-CoV-2-induced lung pathology. Lung sections of animals collected at 7 dpi with SARS-CoV-2 showed widespread inflammation characterized by a cellular infiltrate and airspace consolidation in control protein-treated but not LCB1-Fc treated or naïve mice (**Fig 2B**). At 4 dpi, inflammatory cytokine and chemokine RNA signatures in the lung were absent in LCB1-Fc treated but not control binder treated animals, suggesting that LCB1-Fc treatment prevents virus infection and inflammation in the lung (**Fig 2C and S1**).

**Figure 2.**
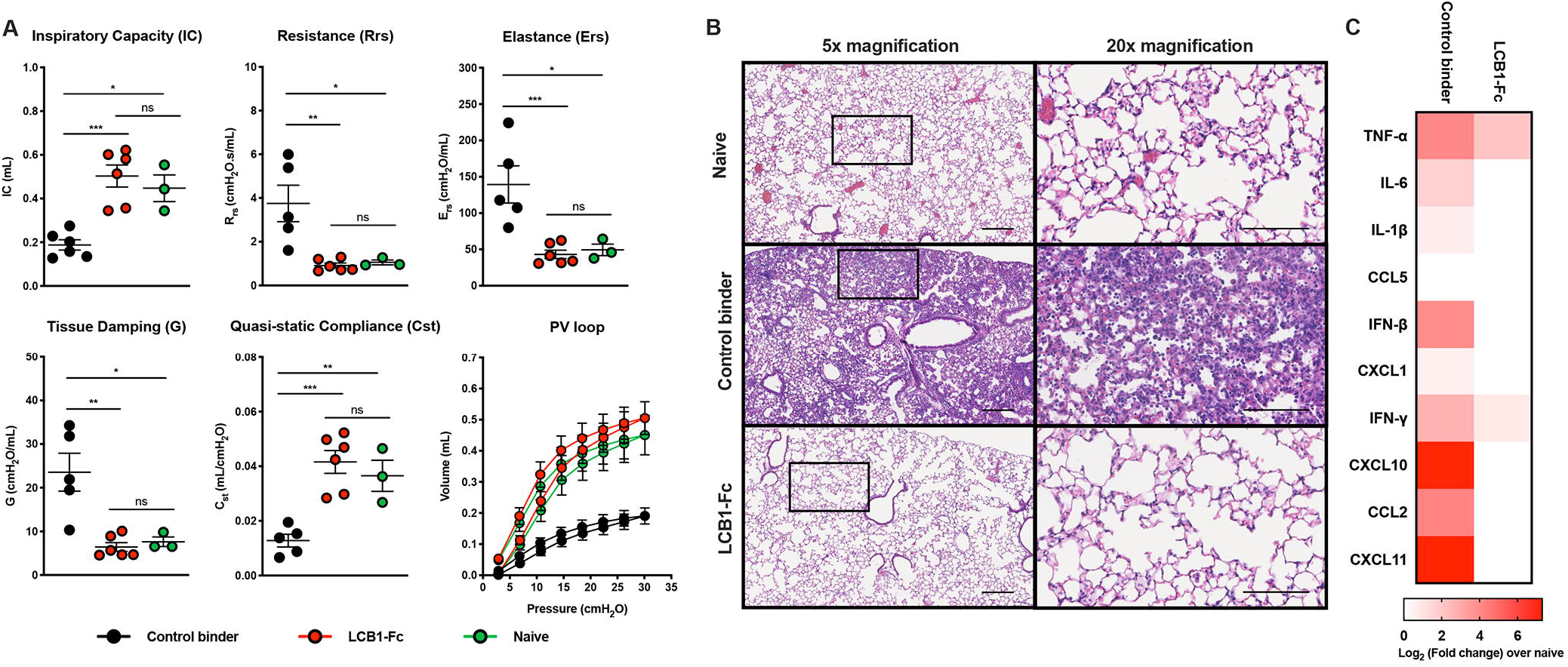
LCB1-Fc prophylaxis prevents SARS-CoV-2-mediated lung disease. (**A**) Respiratory mechanics parameters: inspiratory capacity, resistance, elastance tissue damping, quasi-static compliance, and pressure-volume loops measured at 7 dpi (n = 3-6, two experiments: two-way ANOVA with Tukey’s post-test: ns, not significant, * *P* < 0.05, ** *P* < 0.01, *** *P* < 0.001 between indicated groups). (**B**) Hematoxylin and eosin staining of lung sections from mice treated at D-1 and collected at 7 dpi with SARS-CoV-2. Images show low (left) and high (right; boxed region from left) magnification. Scale bars for all images, 100 μm. Representative images from n = 3 mice per group. (**C**) Heat-map of cytokine mRNA levels from lung tissues of SARS-CoV-2 infected mice at 4 dpi. For each cytokine, the fold-change was calculated relative to age-matched naïve control animals after normalization to *Gapdh* and the Log_2_(fold change) was plotted (n = 8 mice/group relative to n = 3 naïve controls).

### Post-exposure therapy with anti-RBD binders reduces viral burden

To evaluate its efficacy in a post-exposure setting, we administered LCB1-Fc by i.p. injection at 1 dpi. Therapy with LCB1-Fc prevented weight loss (**Fig 3A**) and reduced viral burden in all tested tissues at 4 and 7 dpi (**Fig 3B-G**). Infectious virus was not recovered from the lungs of LCB1-Fc treated animals collected at either timepoint. Lung sections confirmed that therapy with LCB1-Fc improved pathological outcome (**Fig 3H**). At 7 dpi, immune cell infiltrates were absent in the lung sections of LCB1-Fc treated but not control binder-treated animals.

**Figure 3.**
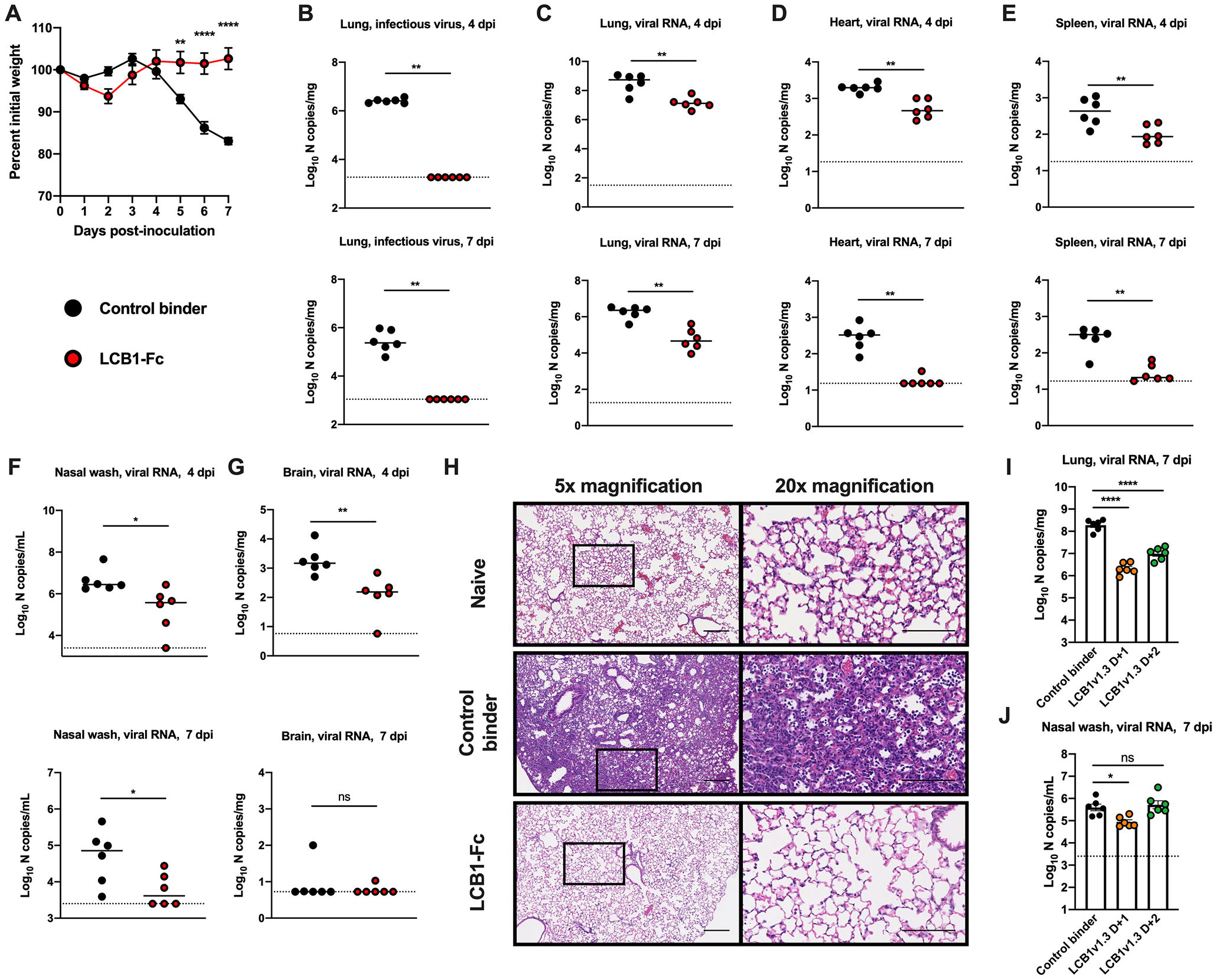
Post-exposure delivery of anti-RBD binders reduces SARS-CoV-2 burden. (**A-G**) 7 to 8-week-old female and male K18-hACE2 transgenic mice received 250 μg of LCB1-Fc or control binder by i.p. injection one day after i.n. inoculation with 10^3^ PFU of SARS-CoV-2. Tissues were collected at 4 or 7 dpi. (**A**) Weight change following LCB1-Fc administration (mean ± SEM; n = 6, two experiments: two-way ANOVA with Sidak’s post-test: ** *P* < 0.01, **** *P* < 0.0001). (**B**) Infectious virus in the lung measured by plaque assay at 4 or 7 dpi in the lung (n = 6, two experiments: ** *P* < 0.01). (**C-G**) Viral RNA levels at 4 or 7 dpi in the lung, heart, spleen, brain, or nasal wash (n = 6, two experiments: Mann-Whitney test: ns, not significant, * *P* < 0.05, ** *P* < 0.01). (**H**) Hematoxylin and eosin staining of lung sections from mice treated at D+1 and collected at 7 dpi with SARS-CoV-2. Images show low (left) and high (right; boxed region from left) magnification. Scale bars for all images, 100 μm. Representative images from n = 3 mice per group. (**I-J**) 7 to 8-week-old male K18-hACE2 transgenic mice received a single 50 μg i.n. dose of LCB1v1.3 or control binder at one- or two-days post-inoculation with 10^3^ PFU of SARS-CoV-2. Viral RNA levels at 7 dpi in the lung (**I**) or nasal wash (**J**) (n = 6, two experiments: one-way ANOVA: ns, not significant, * *P* < 0.05, **** *P* < 0.0001).

We next tested the efficacy of LCB1v1.3 as an i.n.-delivered post-exposure therapy. I.n. delivery, might enable self-administration of an anti-SARS-CoV-2 biological drug. Indeed, miniprotein inhibitors against influenza virus have shown efficacy as a nasal mist (Chevalier et al., 2017). For these studies, we used LCB1v1.3 because it can bind an increased number of RBD molecules for a given mass dose, resulting in increased neutralization activity (**Fig 1C**). Whereas high levels of SARS-CoV-2 RNA were detected in the lungs and other peripheral tissues of control binder-treated animals at 7 dpi, infection was reduced in animals receiving LCB1v1.3 by i.n. administration at D+1 or D+2 after inoculation with SARS-CoV-2 (**Fig 3I and S2**). Levels of viral RNA were reduced in the nasal washes of animals receiving LCB1v1.3 after treatment at D+1 but not D+2 compared to control binder-treated animals (**Fig 3J**).

### Intranasal delivery of LCB1v1.3 confers protection against SARS-CoV-2 when administered up to 5 days before infection

We next evaluated the durability of LCB1v1.3 administered via i.n. prophylaxis. At 5 days, 3 days, 1 day, or 6 hours prior to inoculation with 10^3^ PFU of SARS-CoV-2, K18-hACE2 transgenic mice received a single 50 μg i.n. dose of LCB1v1.3 or the control binder. At 4 or 7 dpi, viral burden in tissues was determined by RT-qPCR. As expected, protection by LCB1v1.3 was better when administered closer to the time of SARS-CoV-2 exposure, as reflected by greater reductions in viral load and weight loss (**Fig 4A-D and S3**). However, even mice receiving LCB1v1.3 five days prior to inoculation and collected at 7 dpi showed reduced viral RNA levels in the lung compared to control binder treated animals. Regardless of the collection timepoint, lung viral RNA levels were reduced in animals receiving LCB1v1.3 three days prior to inoculation with SARS-CoV-2.

**Figure 4.**
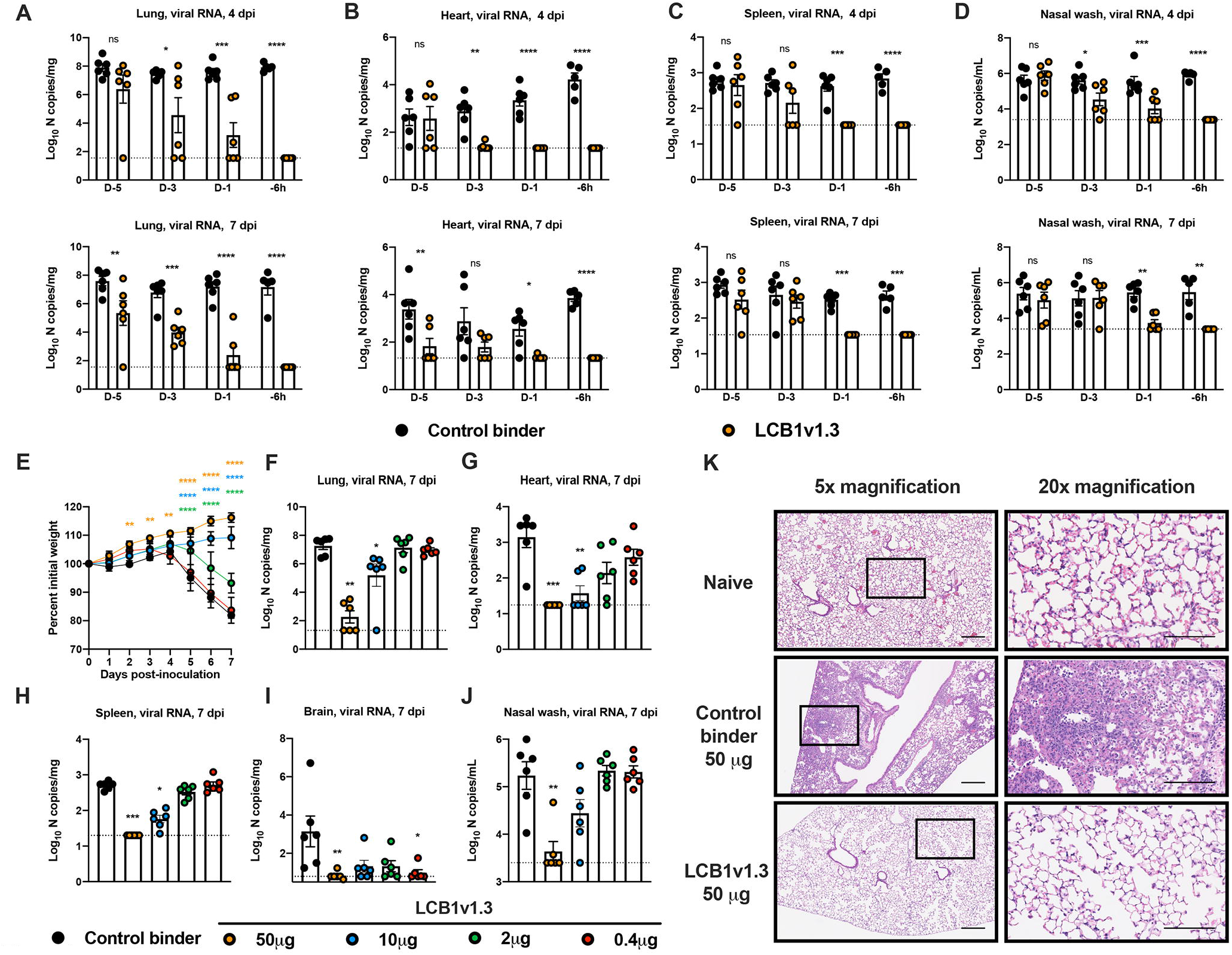
Intranasal administration of LCB1v1.3 reduces viral infection even when given 5 days prior to SARS-CoV-2 exposure. (**A-D**) 7 to 8-week-old female K18-hACE2 transgenic mice received a single i.n. 50 μg dose of LCB1v1.3 or control binder at the indicated time prior to i.n. inoculation with 10^3^ PFU of SARS-CoV-2. Tissues were collected at 4 or 7 dpi and viral RNA levels were determined (n = 5-6 animals per group, two-experiments: two-way ANOVA with Sidak’s post-test: ns, not significant, * *P* < 0.05, ** *P* < 0.01, *** *P* < 0.001, **** *P* < 0.0001). (**E-J**) 7 to 8-week-old female K18-hACE2 transgenic mice received the indicated i.n. dose of LCB1v1.3 or control binder at one day prior to i.n. inoculation with 10^3^ PFU of SARS-CoV-2. (**E**) Weight change following LCB1v1.3 or control binder administration (mean ± SEM; n = 6, two experiments: two-way ANOVA with Sidak’s post-test compared to the control binder treated group: ** *P* < 0.01, **** *P* < 0.0001). (**F-J**) Viral RNA levels at 7 dpi in the lung, heart, spleen, brain, or nasal wash (n = 6, two experiments: Kruskal-Wallis ANOVA with Dunn’s post-test: * *P* < 0.05, ** *P* < 0.01, *** *P* < 0.001). (**K**) Hematoxylin and eosin staining of lung sections from mice treated with a single i.n. 50 μg dose of LCB1v1.3 or control binder at D-1 and collected at 7 dpi with SARS-CoV-2. Images show low (left) and high (right; boxed region from left) magnification. Scale bars for all images, 100 μm. Representative images from n = 3 mice per group.

An important consideration for our binders as a potential therapy is scalable production and feasible dosing. To begin to address this issue, we tested a range of i.n. doses LCB1v1.3 for efficacy (**Fig 4E-J**). Treatment with as little as 2 μg (0.1 mg/kg) of LCB1v1.3 prevented SARS-CoV-2-induced weight loss. Doses between 2 and 10 μg (0.1 to 0.5 mg/kg) of LCB1v1.3 reduced viral RNA levels in the lung, heart, and spleen at 7 dpi relative to control binder-treated animals. Moreover, animals receiving a 50 μg dose of LCB1v1.3 showed minimal, if any, lung inflammation (**Fig 4K**). Collectively, these results indicate that even low doses of LCB1v1.3, when administered via an i.n. route prior to exposure, can limit SARS-CoV-2 infection and disease in the stringent K18-hACE transgenic mouse model of pathogenesis.

### LCB1v1.3 is weakly immunogenic and retains protective activity after repeated dosing

One concern for biological drugs is their potential immunogenicity, which could limit bioavailability and efficacy. To begin to address this concern, we treated K18-hACE2 transgenic mice with 50 μg of control binder or LCB1v1.3 every three days for a total of 18 days (**Fig 5A**). At this time, we collected sera and assessed the presence of anti-LCB1v1.3 antibodies. Only 1 of 10 mice developed IgG antibodies against LCB1v1.3 (**Fig 5B**). To determine if repeated dosing affected LCB1v1.3-mediated protection, we challenged the cohort with 10^3^ PFU of SARS-CoV-2. Again, substantial protection against weight loss (**Fig 5C**) and viral infection in the lung and other organs was observed in all animals receiving LCB1v1.3 (**Fig 5D-H**).

**Figure 5.**
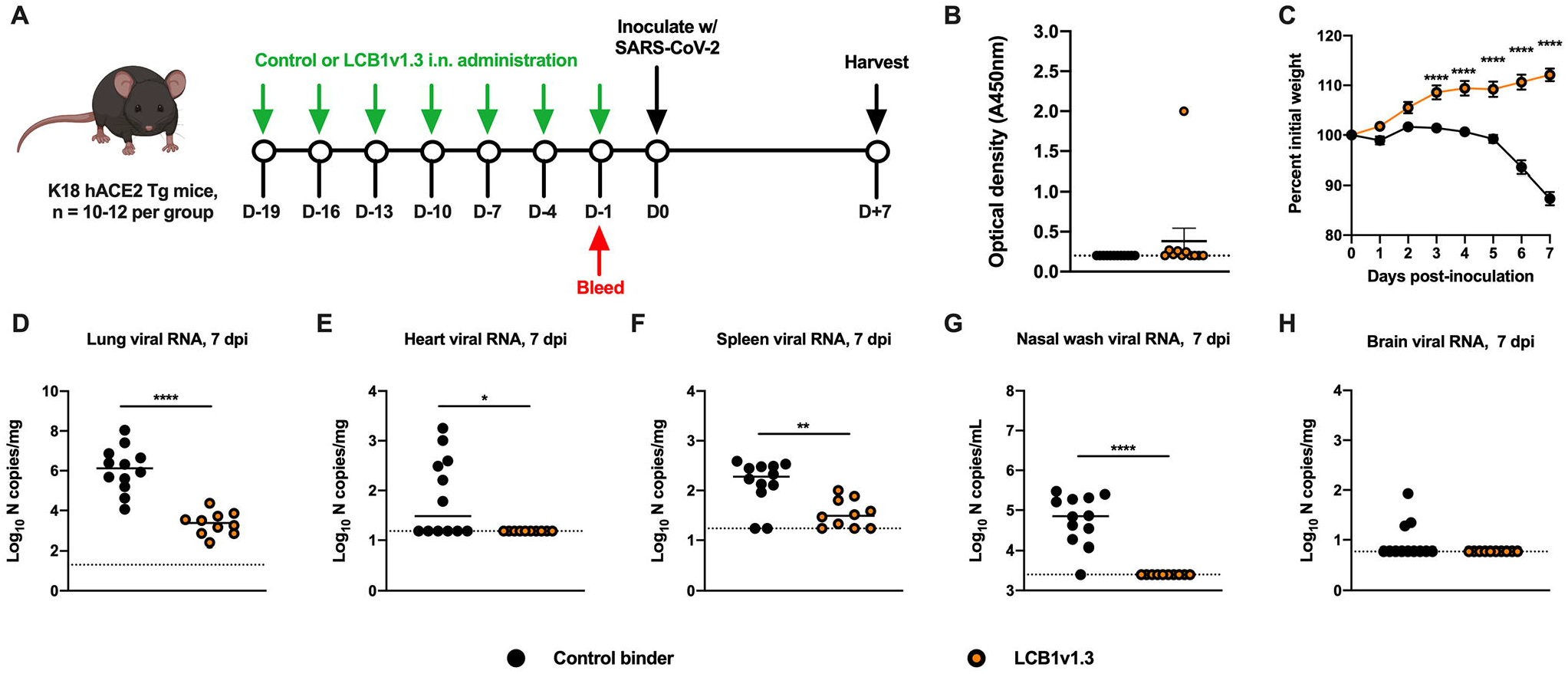
Immunogenicity of LCB1v1.3 and protection from challenge. (**A**) Scheme of experimental details. K18-hACE2 transgenic mice (n = 10 to 12 per group) were treated every 3 days with 50 μg of LCB1v1.3 or control binder by i.n. administration. On day 18 post-treatment, animals were bled and anti-LCB1v1.3 antibodies were measured. The following day, animals were challenged with 10^3^ PFU of SARS-CoV-2, and tissues were collected at 7 dpi. (**B**) Binding of serum antibodies to LCB1v1.3 as measured by ELISA (three experiments). Dashed line indicated limit of detection of the assay. (**C**) Weight change following LCB1v1.3 or control binder administration (mean ± SEM; two experiments: two-way ANOVA with Sidak’s post-test: **** *P* < 0.0001). (**D-H**) Viral RNA levels at 7 dpi in the lung, heart, spleen, brain, or nasal wash (two experiments: Mann-Whitney test: * *P* < 0.05, ** *P* < 0.01, **** *P* < 0.0001).

### LCB1v1.3 protects against emerging SARS-CoV-2 variants

The emergence of variant strains harboring possible escape mutations is of great concern for antibody-based countermeasures that were designed against historical SARS-CoV-2 spike proteins (Chen et al., 2021; Wang et al., 2021a; Wang et al., 2021b; Wibmer et al., 2021). Accordingly, we evaluated the activity of LCB1v1.3 against a B.1.1.7 isolate containing deletions at 69-70 and 144-145, and substitutions at N501Y, A570D, D614G, and P681H, and against a recombinant WA1/2020 strain containing key substitutions present in the B.1.351 and B.1.248 variant strains at residues E484K, N501Y, and D614G (Xie et al., 2021a). Although the neutralizing activity of LCB1v1.3 against the B.1.1.7 and E484K/N501Y/D614G strains was approximately 45 to 50-fold lower than for the WA1/2020 strain, the EC_50_ values still were ~800 pM and 667 pM, respectively (**Fig 6A**). To determine whether LCB1v1.3 could protect *in vivo* against SARS-CoV-2 strains with concerning spike protein substitutions, we treated K18-hACE2 transgenic mice with a single i.n. 50 μg dose of LCB1v1.3 or control binder one day prior to inoculation with 10^3^ PFU of B.1.1.7 or E484K/N501/D614G SARS-CoV-2. Notably, LCB1v1.3 treatment before challenge with either variant strain protected against weight loss (**Fig 6B and 6H**) and viral infection in all tissues collected at 6 dpi (**Fig 6C-G and 6I-M**). Thus, LCB1v1.3 is effective against both circulating and emerging strains of SARS-CoV-2.

**Figure 6.**
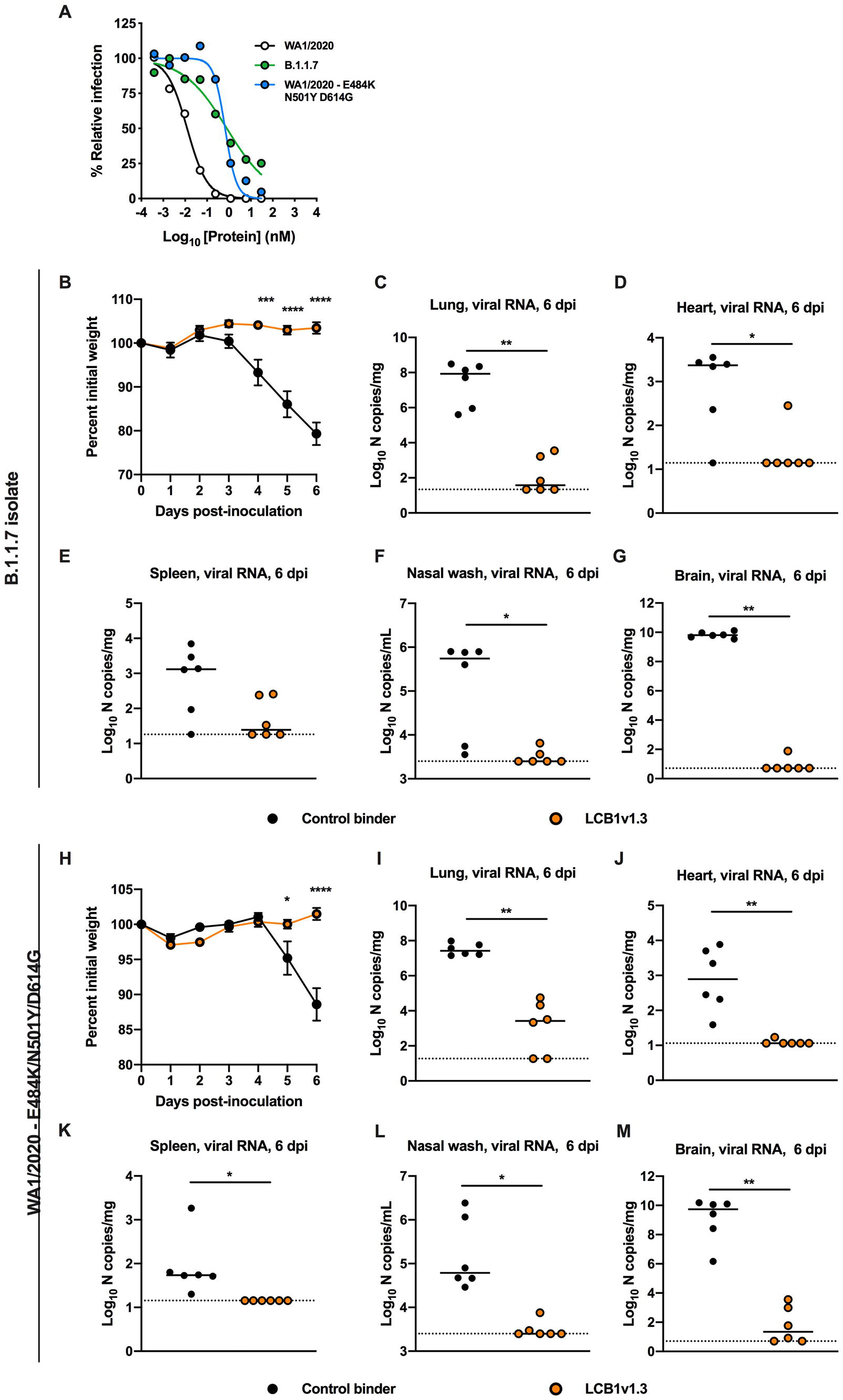
LCB1v1.3 protects mice against B.1.1.7 variant and WA1/2020 E484K/N501Y/D614G strains. (**A**) Neutralization of LCB1v1.3 against B.1.1.7 or WA1/2020 E484K/N501Y/D614G SARS-CoV-2 (EC_50_ values: 802 pM and 667 pM, respectively; mean of two experiments, each performed in duplicate). (**B-G**) 7 to 8-week-old female K18-hACE2 transgenic mice were treated with a single 50 μg i.n. dose of LCB1v1.3 or control binder at 1 day prior to i.n. inoculation with 10^3^ PFU of B.1.1.7. (**B**) Weight change following LCB1v1.3 or control binder administration (mean ± SEM; n = 6, two experiments: two-way ANOVA with Sidak’s post-test: *** *P* < 0.001, **** *P* < 0.0001). (**C-G**) Viral RNA levels at 6 dpi in the lung, heart, spleen, nasal wash, or brain (n = 6, two experiments: Mann-Whitney test: * *P* < 0.05, ** *P* < 0.01). (**H-M**) 8-week-old male K18-hACE2 transgenic mice were treated with a single 50 μg i.n. dose of LCB1v1.3 or control binder at 1 day prior to i.n. inoculation with 10^3^ PFU of WA1/2020 E484K/N501Y/D614G. (**H**) Weight change following LCB1v1.3 or control binder administration (mean ± SEM; n = 6, two experiments: two-way ANOVA with Sidak’s post-test: * *P* < 0.05, **** *P* < 0.0001). (**I-M**) Viral RNA levels at 6 dpi in the lung, heart, spleen, nasal wash, or brain (n = 6, two experiments: Mann-Whitney test: * *P* < 0.05, ** *P* < 0.01).

## DISCUSSION

Here, using the stringent K18-hACE2 mouse model of SARS-CoV-2 pathogenesis, we show that LCB1-Fc, an Fc-containing version of a previously reported SARS-CoV-2 RBD binding miniprotein, LCB1 (Cao et al., 2020), prevented SARS-CoV-2 infection and disease when administered one day before or after virus inoculation. Lung biomechanics of mice treated with LCB1-Fc mirrored those of naïve animals in all parameters tested. Although formal pharmacokinetics studies are needed to establish half-life and bioavailability of our miniprotein, we expect the addition of the Fc moiety improves systemic levels. While our protection studies with LCB1v1.3, which lacks an Fc domain, suggest that our miniprotein binders confer protection through their neutralizing activity, engagement of complement and FcγRs could augment the therapeutic activity of LCB1-Fc, as described recently with anti-SARS-CoV-2 monoclonal antibodies (Schafer et al., 2021; Winkler et al., 2020b). These functions could be explored using loss-of-Fc function (*e.g*., LALA-PG) variants of LCB1-Fc. Depending on the outcome, Fc effector functions could be optimized further through glycan modification or Fc mutation (Kang and Jung, 2019).

We also evaluated the efficacy of LCB1v1.3, an optimized, monomeric form of LCB1 without an Fc domain. A single i.n. dose of LCB1v1.3 reduced viral burden when administered as many as five days before or two days after SARS-CoV-2 infection. While several antibody-based intravenous therapies have been developed against SARS-CoV-2, our i.n. delivery approach is unique. I.n. therapy of SARS-CoV-2 has been reported only with type I interferon in a hamster model of disease (Hoagland et al., 2021) and efficacy was limited. The K18-hACE2 mouse model recapitulates several aspects of severe COVID-19, including lung inflammation and reduced pulmonary function (Golden et al., 2020; Winkler et al., 2020a). Importantly, LCB1v1.3 binder treatment before or after infection limited immune cell infiltration and lung inflammation, which prevented tissue damage and compromise of respiratory function. Since K18-hACE2 mice are highly vulnerable to infection, the therapeutic window of treatment is limited (Winkler et al., 2020b). While we observed reductions in lung viral burden from a single i.n. dose of LCB1v1.3 given two days after virus inoculation, it will be important to improve upon this result. Possible ways to achieve this include higher dosing, repeated dosing, or extended half-life engineering. As part of our proof-of-principle studies for a nasal prophylaxis, we observed little immunogenicity of LCB1v1.3, suggesting that repeated dosing may be possible. Evaluation of miniprotein binders in hamsters and NHPs is needed to extend the efficacy data and provide further rationale for human clinical trials.

Although several antibody-based therapies demonstrate promise against SARS-CoV-2, and a few have been granted EUA status, viral evolution could jeopardize these interventions as evidenced by the emerging variants in the United Kingdom (B.1.1.7), South Africa (B.1.351), Brazil (B.1.248), and elsewhere. Indeed, we and others have observed that many monoclonal and polyclonal antibodies showed reduced neutralization activity against several of these variant strains (Chen et al., 2021; Wang et al., 2021a; Wang et al., 2021b; Wibmer et al., 2021; Xie et al., 2021b). In comparison, LCB1v1.3 showed efficacy against historical (WA1/2020) and emerging (B.1.1.7 and E484K/N501Y/D614G) SARS-CoV-2 strains. Based on the cryo-EM structure of the parent LCB1 binder in complex with SARS-CoV-2 RBD (Cao et al., 2020), only the N501Y mutation is expected to affect binding. While we observed a decrease in the neutralizing activity of LCB1v1.3 against the emerging variants, EC_50_ values were still less than 800 pM, suggesting substantial potency was retained. Additional optimization of LCB1-Fc- and LCB1v1.3-RBD binding interactions, through computational design and functional validation, could reduce the effects of variant mutations on neutralizing activity. Moreover, the development of binder combinations that target different regions of the spike protein of multiple emerging SARS-CoV-2 variants is planned.

Compared to other potential SARS-CoV-2 antibody-based treatments, miniproteins have several benefits: (a) due to their smaller size, they can bind each protomer of a single trimeric spike, resulting in greater potency for a given dose; (b) they can be manufactured cost-effectively; (c) if warranted, they can be refined to overcome escape by new SARS-CoV-2 variants; and (d) they can be mixed using linker proteins to generate multimerized constructs that limit resistance. In summary, our data highlight the promise of rational protein antiviral design and support the development of LCB1v1.3 and LCB1-Fc as potent SARS-CoV-2-specific countermeasures.

## ACKNOWLEDGEMENTS

This study was supported by NIH grants (R01 AI157155, Al134907, and UL1TR001439) and the Defense Advanced Research Project Agency (HR001117S0019 and HR0011835403 contract FA8750-17-C-0219), The Audacious Project at the Institute for Protein Design (L. Car. and D.B.), funding from E. and W. Schmidt by recommendation of the Schmidt Futures program (L. Car., R.R., I.G., and D.B.), the Open Philanthropy Project Improving Protein Design Fund (D.B.), Bill and Melinda Gates Foundation #OPP1156262 (L.S., L. Car. R.R., and D.B.). J.B.C. is supported by a Helen Hay Whitney Foundation postdoctoral fellowship, E.S.W. is supported by F30 AI152327, N.M.K. is supported by T32 AI007172, and P.-Y.S. is supported by awards from the Sealy and Smith Foundation, the Kleberg Foundation, the John S. Dunn Foundation, the Amon G. Carter Foundation, the Gilson Longenbaugh Foundation, and the Summerfield Robert Foundation. We thank Lisa Kozodoy and Lexi Walls for support in development of ELISA assays, Cassie Ogohara, and Michael Murphy for support in protein production and purification, the Pulmonary Morphology Core at Washington University School of Medicine for tissue sectioning and slide preparation and SCIREQ Inc. for providing the flexiVent pulmonary mechanics research platform and analysis software.

This work is licensed under a Creative Commons Attribution 4.0 International (CC BY 4.0) license, which permits unrestricted use, distribution, and reproduction in any medium, provided the original work is properly cited. To view a copy of this license, visit https://creativecommons.org/licenses/by/4.0. This license does not apply to figures/photos/artwork or other content included in the article that is credited to a third party; obtain authorization from the rights holder before using such material.

## AUTHOR CONTRIBUTIONS

J.B.C., L.S., D.B., and M.S.D. designed the research. J.B.C., R.E.C., E.S.W., S.S., and N.M.K. performed mouse experiments and clinical analyses. J.B.C. and B.Y. performed viral burden analysis. J.B.C. and E.S.W. performed pulmonary mechanics analysis. A.L.B. analyzed the tissue sections for histopathology. J.B.C and R.E.C. performed neutralization analysis. L.Cao optimized protein designs, generated computational models, and performed BLI analysis. L. Car. and R.R. purified and prepared the miniproteins. I.G. and L.S. developed and performed ELISA analysis. X.X. and P.Y.S. provided the recombinant virus strain. J.B.C. and M.S.D. wrote the initial draft, with other authors providing editorial comments and helpful discussions about the research.

## SUPPLEMENTAL FIGURE LEGENDS

**Figure S1. Cytokine and chemokine induction following SARS-CoV-2 infection, Related to Fig 2C.** Individual plots for cytokine and chemokine RNA levels in the lungs of SARS-CoV-2 infected mice at 4 dpi following treatment with control or LCB1-Fc binders (n = 8 per group, two experiments: Mann-Whitney test: ns, not significant, * *P* < 0.05, ** *P* < 0.01, *** *P* < 0.001). Data were used to generate the heat-map in **Figure 2C**.

**Figure S2. Intranasal delivery of LCB1v1.3 at 1 or 2 days post-SARS-CoV-2 infection reduces viral burden, Related to Fig 3.** (**A-C**) 7 to 8-week-old male K18-hACE2 transgenic mice received a single 50 μg i.n. dose of LCB1v1.3 or control binder at one- or two-days post-inoculation with 10^3^ PFU of SARS-CoV-2. Viral RNA levels at 7 dpi in the heart (**A**), spleen (**B**), or brain (**C**) (n = 6, two experiments: one-way ANOVA: * *P* < 0.05, ** *P* < 0.01).

**Figure S3. Intranasal prophylaxis of LCB1v1.3 reduces weight loss, Related to Fig 4.** 7 to 8-week-old female K18-hACE2 transgenic mice received a single 50 μg i.n. dose of LCB1v1.3 or control binder at the indicated time prior to i.n. inoculation with 10^3^ PFU of SARS-CoV-2. Weight change was recorded daily (mean ± SEM; n = 6, two experiments: two-way ANOVA with Sidak’s post-test: * *P* < 0.05, ** *P* < 0.01, *** *P* < 0.001, **** *P* < 0.0001).

## STAR METHODS

### RESOURCE AVAILABLITY

#### Lead Contact

Further information and requests for resources and reagents should be directed to the Lead Contact, Michael S. Diamond.

#### Materials Availability

All requests for resources and reagents should be directed to the Lead Contact author. This includes mice, antibodies, protein minibinders, and viruses. All reagents will be made available on request after completion of a Materials Transfer Agreement.

#### Data and code availability

All data supporting the findings of this study are available within the paper and are available from the corresponding author upon request.

### EXPERIMENTAL MODEL AND SUBJECT DETAILS

#### Cells and viruses

Vero E6 (CRL-1586, American Type Culture Collection (ATCC), Vero CCL81 (ATCC), Vero-furin (Mukherjee et al., 2016), and Vero-hACE2-TMPRSS2 (a gift of A. Creanga and B. Graham, NIH) were cultured at 37°C in Dulbecco’s Modified Eagle medium (DMEM) supplemented with 10% fetal bovine serum (FBS), 10□mM HEPES pH 7.3, 1□mM sodium pyruvate, 1× non-essential amino acids, and 100□U/ml of penicillin–streptomycin. Additionally, Vero-hACE2-TMPRSS2 cells were cultured in the presence of 5 μg/mL puromycin. The WA1/202 (2019n-CoV/USA_WA1/2020) isolate of SARS-CoV-2 was obtained from the US Centers for Disease Control (CDC). The B.1.1.7 and WA1/2020 E484K/N501Y/D614G viruses have been described previously (Chen et al., 2021; Xie et al., 2021a). Infectious stocks were propagated by inoculating Vero CCL81 or Vero-hACE2-TMPRSS2 cells. Supernatant was collected, aliquoted, and stored at −80°C. All work with infectious SARS-CoV-2 was performed in Institutional Biosafety Committee-approved BSL3 and A-BSL3 facilities at Washington University School of Medicine using positive pressure air respirators and protective equipment.

#### Mouse experiments

Animal studies were carried out in accordance with the recommendations in the Guide for the Care and Use of Laboratory Animals of the National Institutes of Health. The protocols were approved by the Institutional Animal Care and Use Committee at the Washington University School of Medicine (assurance number A3381–01). Virus inoculations were performed under anesthesia that was induced and maintained with ketamine hydrochloride and xylazine, and all efforts were made to minimize animal suffering.

Heterozygous K18-hACE c57BL/6J mice (strain: 2B6.Cg-Tg(K18-ACE2)2Prlmn/J) were obtained from The Jackson Laboratory. Animals were housed in groups and fed standard chow diets. Mice of different ages and both sexes were administered 10^3^ PFU of SARS-CoV-2 via intranasal administration.

### METHOD DETAILS

#### Miniprotein production

LCB1-Fc was synthesized and cloned by GenScript into pCMVR plasmid, with kanamycin resistance. Plasmids were transformed into the NEB 5-alpha strain of *E. coli* (New England Biolabs) to recover DNA for transient transfection into Expi293F mammalian cells. Expi293F cells were grown in suspension using Expi293F expression medium (Life Technologies) at 33°C, 70% humidity, and 8% CO_2_ rotating at 150 rpm. The cultures were transfected using PEI-MAX (Polyscience) with cells grown to a density of 3 x 10^6^ cells per mL and cultivated for 3 days. Supernatants were clarified by centrifugation (5 min at 4000 x *g*, addition of PDADMAC solution to a final concentration of 0.0375% (Sigma Aldrich, #409014), and a second spin (5 min at 4000 x *g*). Clarified supernatants were purified using a MabSelect PrismA 2.6×5 cm column (Cytiva) on an AKTA Avant150 FPLC (Cytiva). Bound protein was washed with five column volumes of 20 mM NaPO_4_ and 150 mM NaCl pH 7.2, then five column volumes of 20 mM NaPO_4_ and 1 M NaCl pH 7.4, and eluted with three column volumes of 100 mM glycine at pH 3.0. The eluate was neutralized with 2 M Tris base to a final concentration of 50 mM. SDS-PAGE was used to assess protein purity. The protein was passed through a 0.22 μm filter and stored at 4°C until use.

LCB1v1.3 with polar mutations (4N, 14K, 15T, 17E, 18Q, 27Q, 38Q) relative to the original LCB1 was cloned into a pet29b vector. LCB1v1.3 was expressed in Lemo21(DE3) (NEB) in terrific broth media and grown in 2 L baffled shake flasks. Bacteria were propagated at 37°C to an O.D.600 of ~0.8, and then induced with 1 mM IPTG. Expression temperature was reduced to 18°C, and the cells were shaken for ~16 h. The cells were harvested and lysed using heat treatment and incubated at 80°C for 10 min with stirring. Lysates were clarified by centrifugation at 24,000 × *g* for 30 min and applied to a 2.6×10 cm Ni Sepharose 6 FF column (Cytiva) for purification by IMAC on an AKTA Avant150 FPLC system (Cytiva). Proteins were eluted over a linear gradient of 30 mM to 500 mM imidazole in a buffer of 50 mM Tris pH 8.0 and 500 mM NaCl. Peak fractions were pooled, concentrated in 10 kDa MWCO centrifugal filters (Millipore), sterile filtered (0.22 μm) and applied to either a Superdex 200 Increase 10/300, or HiLoad S200 pg GL SEC column (Cytiva) using 50 mM phosphate pH 7.4, 150 mM NaCl buffer. After size exclusion chromatography, bacterial-derived components were tested to confirm low levels of endotoxin.

#### Biolayer interferometry

Biolayer interferometry data were collected using an Octet RED96 (ForteBio) and processed using the instrument’s integrated software. Briefly, biotinylated RBD (Acro Biosystems) was loaded onto streptavidin-coated biosensors (SA ForteBio) at 20 nM in binding buffer (10 mM HEPES (pH 7.4), 150 mM NaCl, 3 mM EDTA, 0.05% surfactant P20, and 0.5% non-fat dry milk) for 360 s. Analyte proteins (LCB1v1.3 or LCB1-Fc) were diluted from concentrated stocks into binding buffer. After baseline measurement in the binding buffer alone, the binding kinetics were monitored by dipping the biosensors in wells containing the target protein at the indicated concentration (association step) for 3,600 s and then dipping the sensors back into baseline/buffer (dissociation) for 7,200 s.

#### Plaque assay

Vero-furin cells (Mukherjee et al., 2016) were seeded at a density of 2.5×10^5^ cells per well in flat-bottom 12-well tissue culture plates. The following day, medium was removed and replaced with 200 μL of 10-fold serial dilutions of the material to be titrated, diluted in DMEM+2% FBS, and plates incubated at 37°C with rocking at regular intervals. One hour later, 1 mL of methylcellulose overlay was added. Plates were incubated at 37°C for 72 h, then fixed with 4% paraformaldehyde (final concentration) in PBS for 20 min. Fixed cell monolayers were stained with 0.05% (w/v) crystal violet in 20% methanol and washed twice with distilled, deionized water.

#### Measurement of viral burden

Tissues were weighed and homogenized with zirconia beads in a MagNA Lyser instrument (Roche Life Science) in 1,000 μL of DMEM media supplemented with 2% heat-inactivated FBS. Tissue homogenates were clarified by centrifugation at 10,000 rpm for 5 min and stored at −80°C. RNA was extracted using the MagMax mirVana Total RNA isolation kit (Thermo Scientific) on a Kingfisher Flex extraction robot (Thermo Scientific). RNA was reverse transcribed and amplified using the TaqMan RNA-to-CT 1-Step Kit (ThermoFisher). Reverse transcription was carried out at 48°C for 15 min followed by 2 min at 95°C. Amplification was accomplished over 50 cycles as follows: 95°C for 15 s and 60°C for 1 min. Copies of SARS-CoV-2 N gene RNA in samples were determined using a previously published assay (Case et al., 2020; Hassan et al., 2020). Briefly, a TaqMan assay was designed to target a highly conserved region of the N gene (Forward primer: ATGCTGCAATCGTGCTACAA; Reverse primer: GACTGCCGCCTCTGCTC; Probe: /56-FAM/TCAAGGAAC/ZEN/AACATTGCCAA/3IABkFQ/). This region was included in an RNA standard to allow for copy number determination down to 10 copies per reaction. The reaction mixture contained final concentrations of primers and probe of 500 and 100 nM, respectively.

#### Cytokine and chemokine mRNA measurements

RNA was isolated from lung homogenates as described above. cDNA was synthesized from DNAse-treated RNA using the High-Capacity cDNA Reverse Transcription kit (Thermo Scientific) with the addition of RNase inhibitor following the manufacturer’s protocol. Cytokine and chemokine expression was determined using TaqMan Fast Universal PCR master mix (Thermo Scientific) with commercial primers/probe sets specific for *IFN-g* (IDT: Mm.PT.58.41769240), *IL-6* (Mm.PT.58.10005566), *IL-1b* (Mm.PT.58.41616450), *Tnfa* (Mm.PT.58.12575861), *CXCL10* (Mm.PT.58.43575827), *CCL2* (Mm.PT.58.42151692), *CCL5* (Mm.PT.58.43548565), *CXCL11* (Mm.PT.58.10773148.g), *Ifnb* (Mm.PT.58.30132453.g), *CXCL1* (Mm.PT.58.42076891) and results were normalized to *GAPDH* (Mm.PT.39a.1) levels. Fold change was determined using the 2^-ΔΔCt^ method comparing treated mice to naïve controls.

#### Lung Pathology

Animals were euthanized before harvest and fixation of tissues. The left lung was first tied off at the left main bronchus and collected for viral RNA analysis. The right lung was inflated with approximately 1.2 mL of 10% neutral buffered formalin using a 3-mL syringe and catheter inserted into the trachea. Tissues were embedded in paraffin, and sections were stained with hematoxylin and eosin. Slides were scanned using a Hamamatsu NanoZoomer slide scanning system, and images were viewed using NDP view software (ver.1.2.46).

#### Respiratory mechanics

Mice were anesthetized with ketamine/xylazine (100 mg/kg and 10 mg/kg, i.p., respectively). The trachea was isolated via dissection of the neck area and cannulated using an 18-gauge blunt metal cannula (typical resistance of 0.18 cmH_2_O.s/mL), which was secured in place with a nylon suture. The mouse then was connected to the flexiVent computer-controlled piston ventilator (SCIREQ Inc.) via the cannula, which was attached to the FX adaptor Y-tubing. Mechanical ventilation was initiated, and mice were given an additional 100 mg/kg of ketamine and 0.1 mg/mouse of the paralytic pancuronium bromide via intraperitoneal route to prevent breathing against the ventilator and during measurements. Mice were ventilated using default settings for mice, which consisted in a positive end expiratory pressure at 3 cm H_2_O, a 10 mL/kg tidal volume (Vt), a respiratory rate at 150 breaths per minute (bpm), and a fraction of inspired oxygen (FiO_2_) of 0.21 (*i.e*., room air). Respiratory mechanics were assessed using the forced oscillation technique, as previously described (McGovern et al., 2013), using the latest version of the flexiVent operating software (flexiWare v8.1.3). Pressure-volume loops and measurements of inspiratory capacity also were performed.

#### Neutralization assay

Serial dilutions of binder proteins were incubated with 10^2^ focus-forming units (FFU) of SARS-CoV-2 for 1 h at 37°C. Binder-virus complexes were added to Vero E6 (WA1/2020) or Vero-hACE2-TMPRSS2 (B.1.1.7 and WA1/2020 E484K/N501Y/D614G) cell monolayers in 96-well plates and incubated at 37°C for 1 h. Subsequently, cells were overlaid with 1% (w/v) methylcellulose in MEM supplemented with 2% FBS. Plates were harvested 24-30 h later by removing overlays and fixed with 4% PFA in PBS for 20 min at room temperature. Plates were washed and sequentially incubated with an oligoclonal pool of SARS2-2, SARS2-11, SARS2-16, SARS2-31, SARS2-38, SARS2-57, and SARS2-71 anti-spike protein antibodies (Zhou et al., 2021) and HRP-conjugated goat anti-mouse IgG in PBS supplemented with 0.1% saponin and 0.1% bovine serum albumin. SARS-CoV-2-infected cell foci were visualized using TrueBlue peroxidase substrate (KPL) and quantitated on an ImmunoSpot microanalyzer (Cellular Technologies). Data were processed using Prism software (GraphPad Prism 8.0).

#### ELISA

C-terminal biotinylated LCB1.1v3 was immobilized on streptavidin-coated plates (RayBiotech #7C-SCP-1) at 2.5 μg/mL in 100 μL total volume per well and incubated at 4°C overnight. Plates were washed with wash buffer (TBS + 0.1% (w/v) BSA + 0.05% (v/v) Tween20) and blocked with 200 μL/well blocking buffer (TBS + 2% (w/v) BSA + 0.05% (v/v) Tween20) for 1 h at room temperature. Plates were rinsed with wash buffer using 200 μL/well, and 100 μL of 1:100 diluted sera samples in blocking buffer were added to respective wells. For a positive control, Fc-RBD was serially diluted 1:5 starting at 240 ng/mL in 100 μL of blocking buffer. All samples were incubated for 1 h at room temperature. Plates were washed using 200 μL/well of wash buffer. For the serum samples, HRP-conjugated horse anti-mouse IgG antibody (Vector Laboratories #PI-2000-1) was diluted 1:200 in blocking buffer, and 100 μL was incubated in each well at room temperature for 30 min. For the positive control, HRP-conjugated mouse anti-human IgG antibody (Invitrogen #05-4220) was diluted 1:500 in blocking buffer, and 100 μL was incubated in each well at room temperature for 30 min. Plates were rinsed with wash buffer, and 100 μL of TMB (SeraCare) was added to each well for 2 min. The reaction was quenched by adding 100 μL of 1N HCl. Optical densities were determined at 450nm on a Synergy Neo2 plate reader (BioTek Instruments).

### QUANTIFICATION AND STATISTICAL ANALYSIS

Statistical significance was assigned when *P* values were < 0.05 using Prism Version 8 (GraphPad). Tests, number of animals, median values, and statistical comparison groups are indicated in each of the Figure legends. Analysis of weight change was determined by two-way ANOVA. Changes in functional parameters or immune parameters were compared to control binder-treated animals and analyzed by one-way ANOVA with multiple comparisons tests. Statistical analyses of viral burden between two groups were determined by Mann-Whitney test.

