## Supplementary figures and images for "Ultrapotent miniproteins targeting the receptor-binding domain protect against SARS-CoV-2 infection and disease in mice"

### Supplemental Figure 1

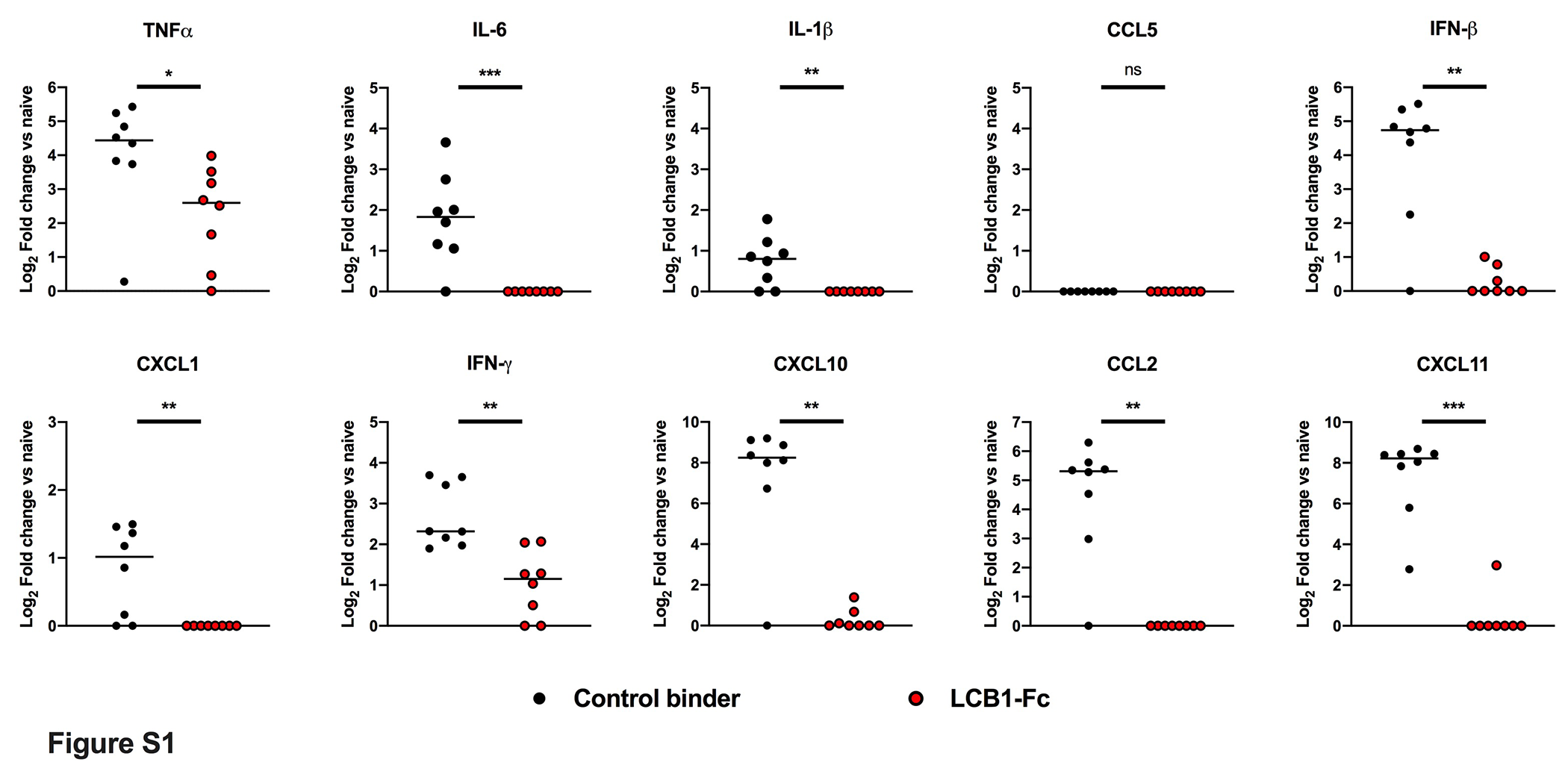

### Supplemental Figure 2

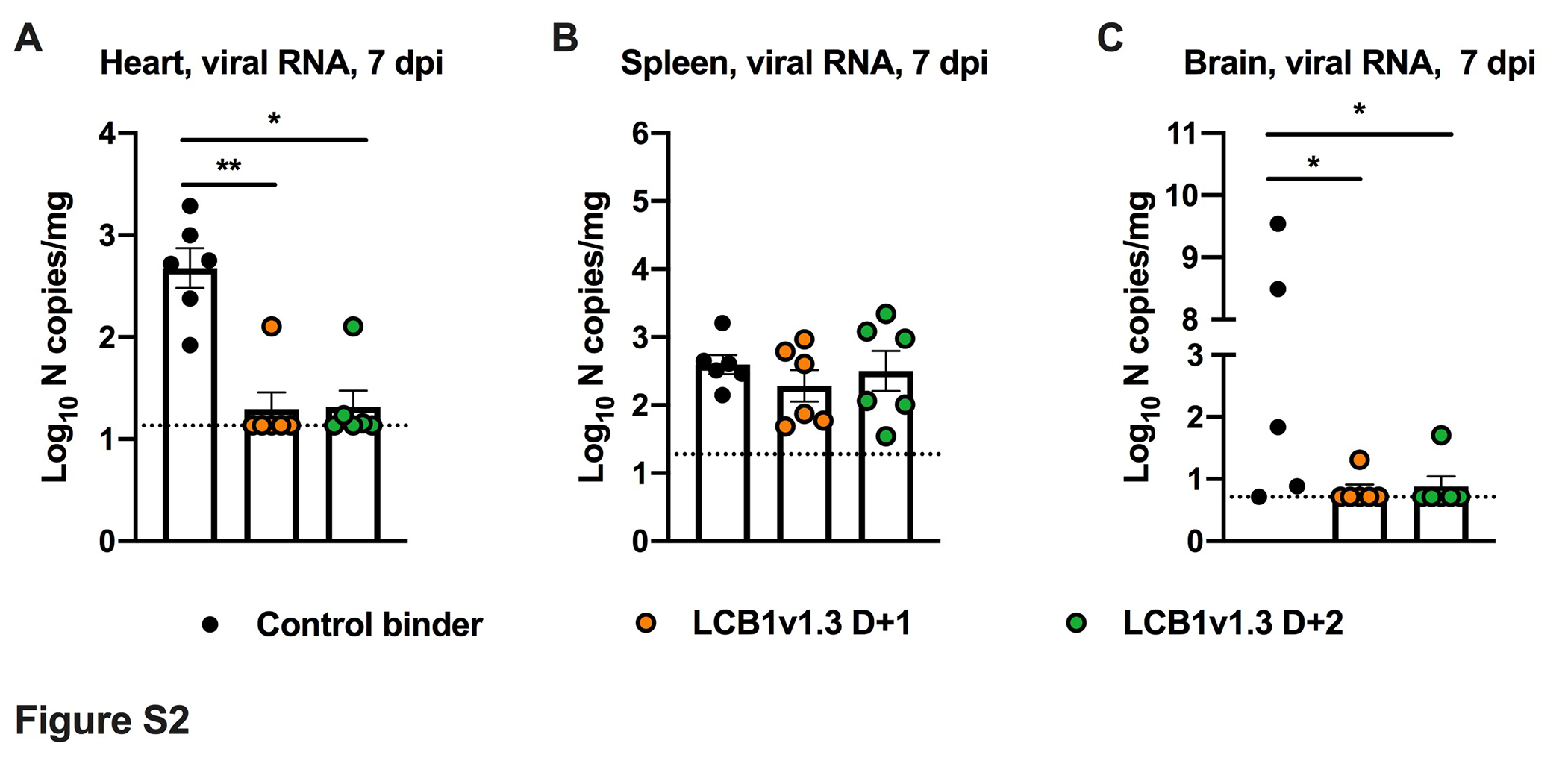

### Supplemental Figure 3

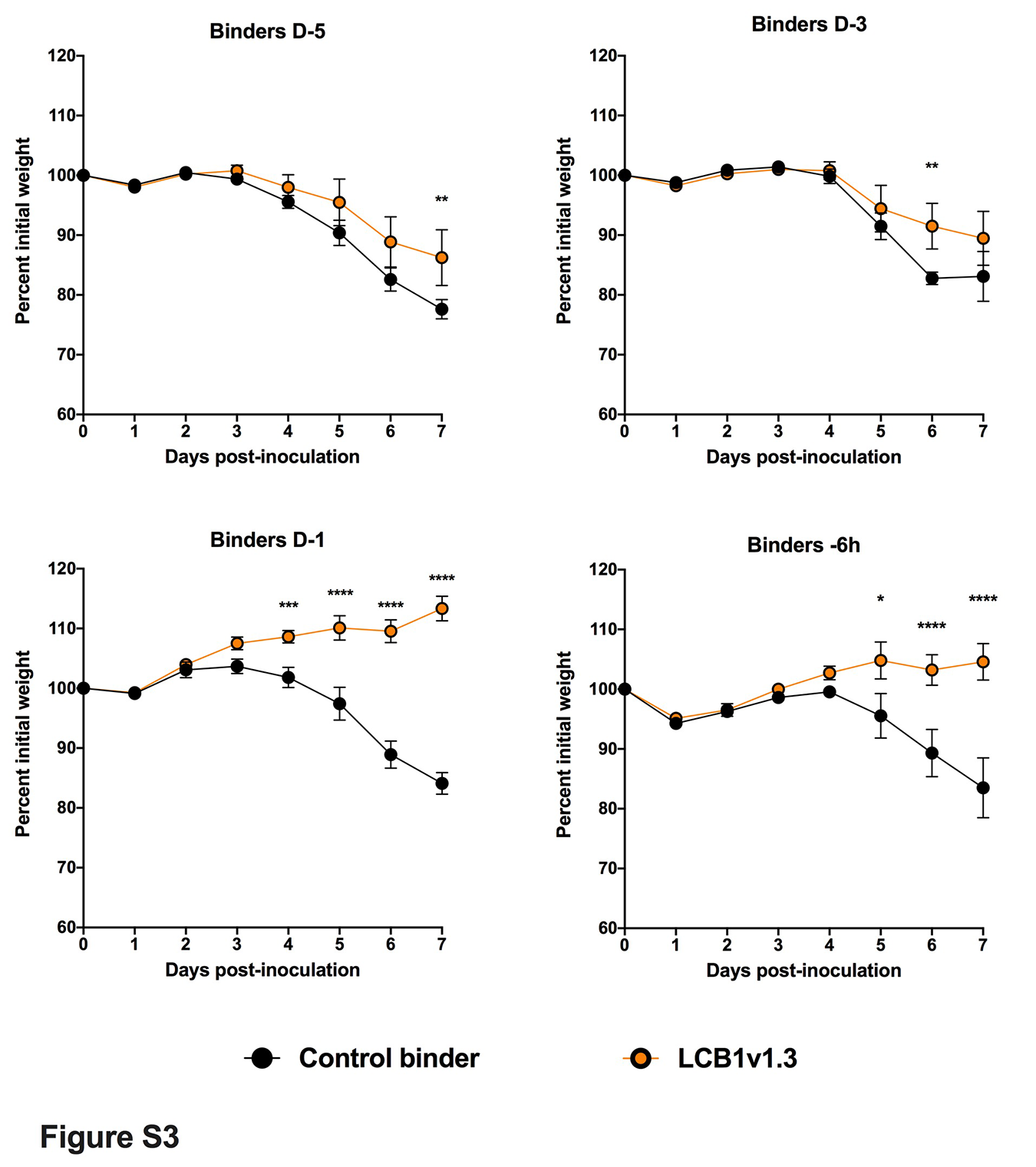
